# Short chain fatty acids regulate the chromatin landscape and distinct gene expression changes in human colorectal cancer cells

**DOI:** 10.1101/2025.05.07.652677

**Authors:** Tohfa Kabir, Charlotte A. Connamacher, Zara Nadeem, Matthew R. Paul, Matthew R. Smutny, Zoe K. Lawler, Annaelle M. Djomo, Thomas S. Carroll, Leah A. Gates

**Author notes:** These authors contributed equally (Tohfa Kabir, Charlotte A. Connamacher).

## Abstract

Short chain fatty acids (SCFAs) are small metabolites that are produced through the activity of microbes and have important roles in human physiology. These metabolites have varied mechanisms in interacting with the host, of which one such mode is decorating the chromatin landscape. We previously detected specific histone modifications in the mouse gut that can be derived from SCFAs and are regulated by the microbiota. However, the roles of these SCFAs on chromatin and how they impact gene regulation in human cells is largely unknown. Now, our studies demonstrate these microbiota-dependent histone posttranslational modifications (PTMs) are associated with alterations in gene regulation in human cells. We show that histone butyrylation on H3K27 is detected in human colon samples. Furthermore, histone acetylation, butyrylation, and propionylation on H3K9 and H3K27 are responsive to levels of SCFAs in human colon cancer cell lines and are associated with active gene regulatory elements. In addition, treatment of human cancer cell lines with individual metabolites or combinations of SCFAs replicating the intestinal lumen environment result in distinct and overlapping gene program changes, with butyrate largely driving gene regulatory effects of SCFA combinations. Lastly, we define butyrate effects on gene expression that are independent of HDAC inhibition and are dependent on p300/CBP, suggesting potential gene programs regulated by histone butyrylation. Together, these results demonstrate that SCFAs are key regulators of the chromatin landscape and gene programs in human colorectal cancer cells.

## Introduction

In the gut, commensal microbes ferment fiber resulting in the accumulation of high concentrations of the metabolites acetate, butyrate, and propionate in the intestinal lumen^1,2^. These SCFAs act on the host epithelium through several different mechanisms or “fates”: for example, they can bind to cell surface or nuclear receptors to induce signaling cascades and specific gene programs, and they can feed into host cell energy pathways. Furthermore, SCFAs can impact the chromatin landscape both directly through serving as donor molecules for histone posttranslational modifications and indirectly by inhibiting select chromatin-modifying enzymes, such as histone deacetylases (HDACs)^3–11^. SCFAs regulate multiple physiological systems including neuronal health, intestinal inflammation, tumor development and metabolic processes such as insulin sensitivity and intestinal gluconeogenesis^3^. It is likely that a combination of molecular mechanisms occurs simultaneously *in vivo* for SCFA action on host physiology. We aimed to investigate the role of SCFA addition to chromatin in gene regulation.

The chromatin landscape is a key regulator of gene transcription, chromatin accessibility, and genomic organization^12,13^. Posttranslational modifications on histone proteins, or histone ‘marks’, allow for the dynamic regulation of chromatin and demarcate genomic regions. Histone marks are composed of diverse chemical moieties, of which many ‘newer’ histone marks have recently been discovered^14^. Since the source of many of these histone marks are small metabolites, metabolite availability is often linked to the global levels of different histone marks^15–18^. Enzymes that deposit histone marks, such as histone acyltransferases (HATs), utilize small metabolites as donor molecules^19^. After coenzyme-A addition, the SCFAs acetate, butyrate, and propionate can be utilized to deposit histone acetylation, butyrylation, and propionylation, respectively^11,20,21^. These SCFA-derived histone marks are part of a class of modifications called histone acyl marks, which are often associated with gene activation and expression^17^.

We recently discovered that in addition to histone acetylation, histone butyrylation and propionylation are localized in the mouse intestinal epithelium and are regulated by the microbiome and metabolite availability^11^. In addition, murine histone butyrylation is associated with high levels of gene expression^11^. Histone propionylation on histone H3 lysine 14 is associated with gene activation in the mouse liver and butyrylation on histone H4 lysine 5 and 8 activates transcription^21,22^. Furthermore, different site-specific marks of histone butyrylation and propionylation were discovered to have distinct roles in human colon cell lines and mouse intestinal cells, which are correlated with open regions of chromatin and gene expression^23^. Here, we aimed to define how histone butyrylation and propionylation sites found in the mouse intestine act in human cells in response to SCFA signaling. In addition, we focus the “fate” of butyrate as histone butyrylation using a targeted approach in human colorectal cancer cells.

## Results

To investigate the role of different histone acyl marks in human colon cells, we first wanted to establish whether these modifications are detectable in human tissue. We therefore stained human intestinal sections for one representative non-acetyl histone acyl marks that we previously detected in the mouse gut: histone H3 butyrylation on lysine 27 (H3K27bu). This histone mark was readily detected by immunofluorescence staining in colon sections from two different patients, while we detected minimal staining in the neighboring ileum (**Figure 1a**). This data suggests that histone butyrylation is present in the human colon.

**Figure 1:**
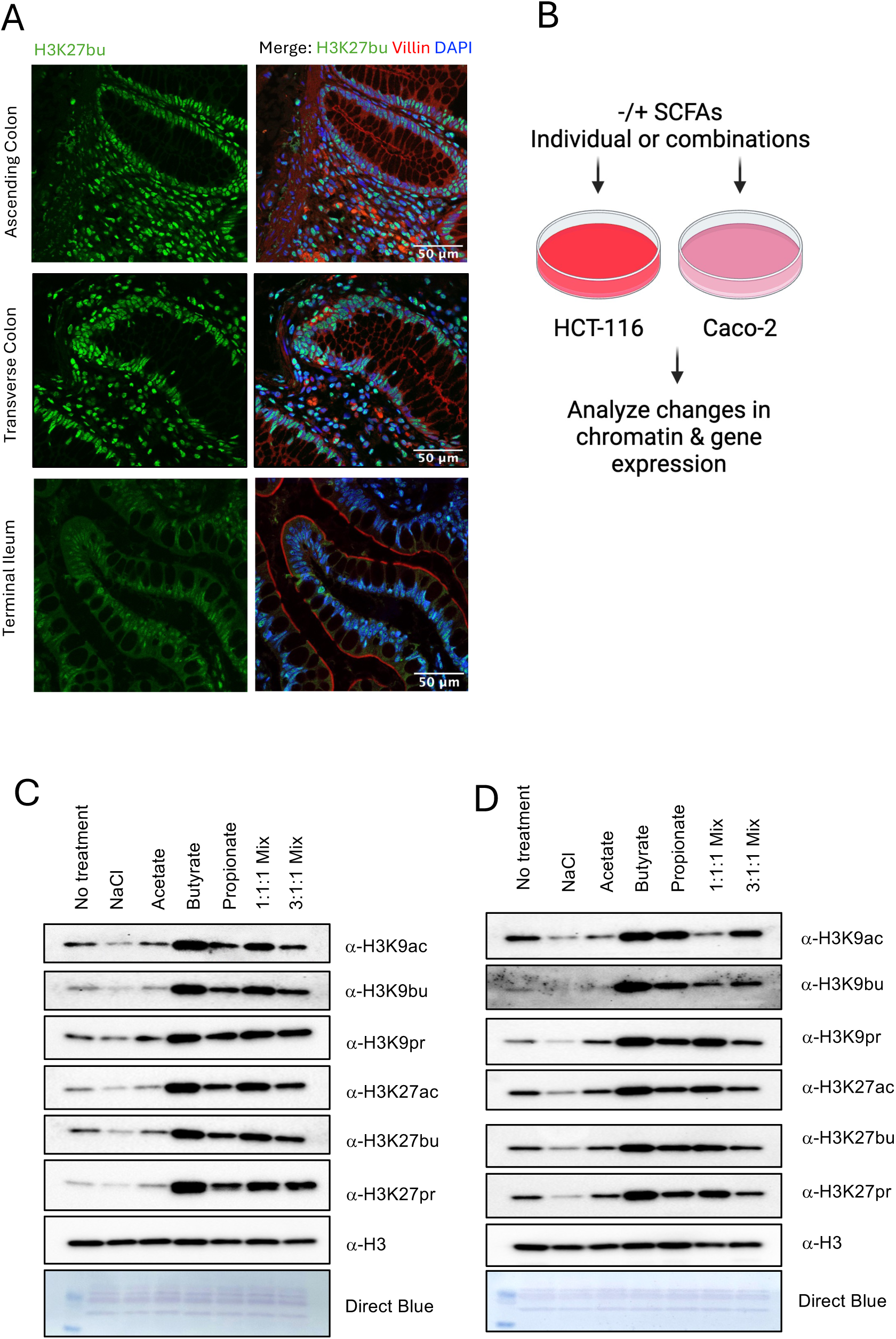
Histone acylations are detected in human colon cells and regulated by metabolites. (A) Histone butyrylation on H3 lysine 27 is detected in human colon samples. Representative immunofluorescence images of H3K27bu, Villin (marker of epithelium), and DAPI in human colon and ileum sections. The colon images are of sections from two different patients. Scale bar = 50 μm. (B) Schematic for cell culture treatments with SCFAs. Image was created with Biorender.com (C-D) Select SCFA treatment results in global increases of histone acyl marks. Representative immunoblots of n =3 independent biological replicates from (C) HCT-116 cells grown in HPLM and (D) Caco-2 cells grown in standard EMEM growth media. Cells were treated with a sodium matched control (NaCl), 5 mM of single SCFAs, an equimolar combination (1.67 mM) each of acetate, butyrate, and propionate (1:1:1 mix), or physiological combination (3:1:1 mix of 3 mM acetate, 1 mM butyrate, 1 mM propionate) for 24 hours. NaCl serves as a control for sodium addition and H3 serves as a loading control.

We then aimed to investigate how the exogenous treatment of SCFAs regulate different histone marks by treating human colon cancer cell lines, HCT-116 and Caco-2, with different types of SCFAs. We reasoned that the addition of SCFAs may result in the deposition of SCFA-derived histone marks onto chromatin. We decided to focus on the same histone marks that we had previously identified *in vivo* in mouse intestinal epithelial cells: acetylation, butyrylation, and propionylation on histone H3 lysine 9 and 27^11^. Treatment with butyrate, propionate, or a mixture of all three SCFAs resulted in a global increase in histone butyrylation or propionylation **(Figure 1b-d, Supplemental Figure 1-2)**. While these modifications display similar patterns, our previous work demonstrates that these antibodies are specific to their respective histone marks^11^. In addition, histone acetylation was increased following butyrate and propionate treatments but was not affected by acetate treatment. This was expected, since typically increases in histone acetylation following exogenous acetate treatment is only observed when cells are grown in low glucose^24^. Together, these data suggest that global levels of histone acyl marks on H3K9 and H3K27 are increased following butyrate or propionate treatments in human colon cancer cell lines.

We then mapped the genomic localization of the different histone acyl marks. Chromatin immunoprecipitation followed by sequencing (ChIP-seq) revealed that all histone acyl marks display similar patterns of localization (**Figure 2a-b**). H3K9bu had a weaker occupancy signal, but still followed the same basic pattern observed for the other marks. Based on the tracks and similar localization to other active histone marks like H3K4me3, it seemed that all of the histone acyl marks interrogated overlapped with active genomic regions or open chromatin. We further analyzed the localization of sequencing reads, which revealed increased proportions of acyl mark ChIP-seq reads in gene promoter regions, 5’ untranslated regions, and coding sequences (**Figure 2c**). These observations suggest that the histone acyl marks all have similar localization patterns that overlap with active chromatin and gene regulatory elements, which is consistent with other reports^21–23^.

**Figure 2:**
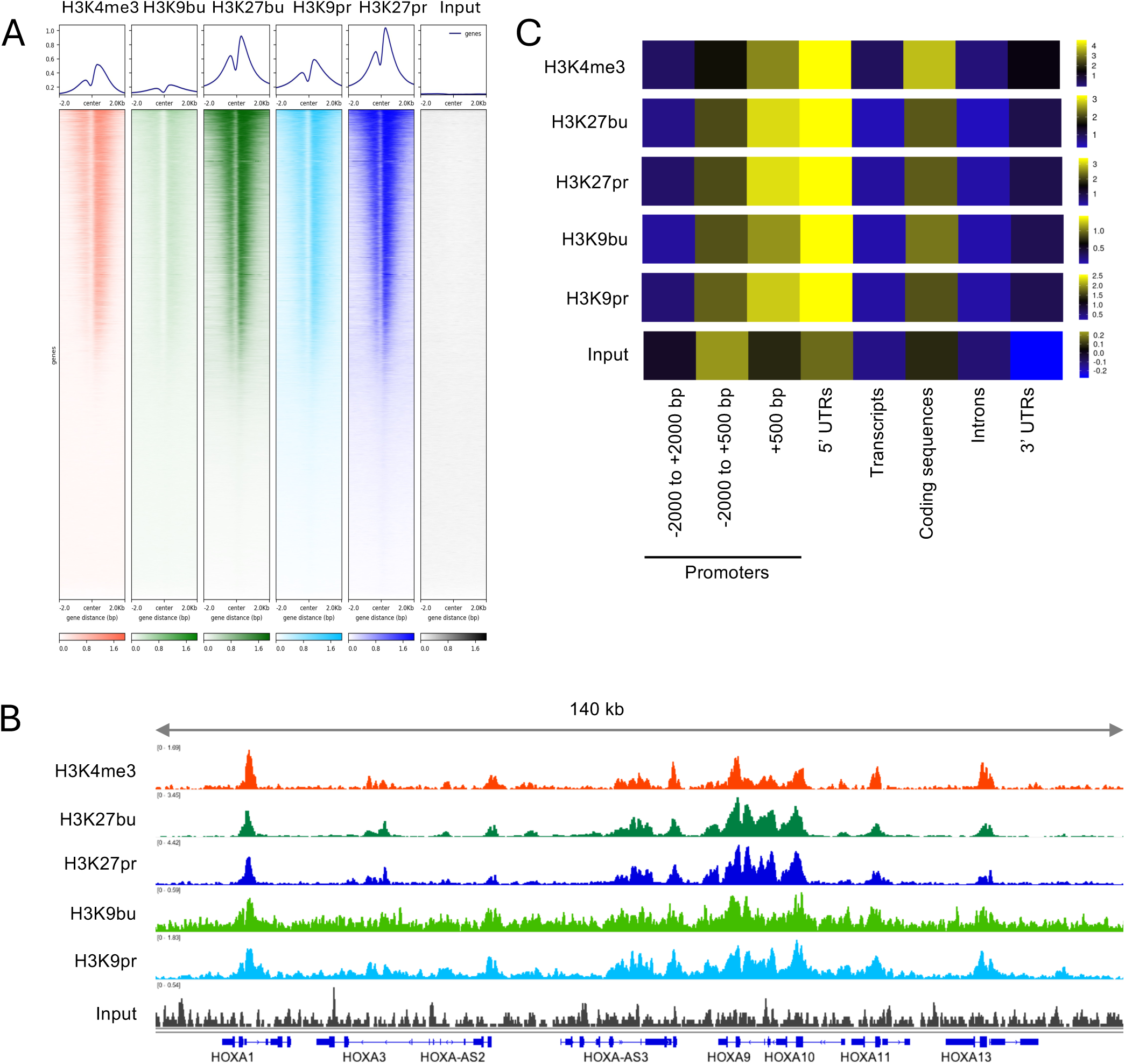
Histone propionylation and butyrylation are associated with open chromatin in human colon cancer cells. (A) Global localization of different histone acyl marks in HCT-116 cells. ChIP-seq was performed in HCT-116 cells grown in DMEM and treated with different SCFAs for 24 hours. Heatmaps of ChIP-seq data depicting localization of different histone acyl marks and a positive control, H3K4me3 at TSS +/- 2kb. Heatmaps were generated using deepTools. (B) Genomic distribution of ChIP-seq reads. Enrichment of reads within the listed annotated regions are displayed as a heatmap. The enrichment is normalized to the proportion of these elements in the genome. Data was generated with ChIPQC and representative ChIPs are shown. (C) Representative tracks depicting ChIP-seq data. Tracks represent normalized, fragment-extended signal bigWigs. All tracks are individually autoscaled and figure was generated using IGV.

Next, we interrogated how exogenous SCFA treatment regulates gene expression. We first treated HCT-116 cells with either single SCFAs or a combination of all three SCFAs where all treated cells received equimolar treatments. We chose a six-hour time point to better capture potential direct gene expression changes. We observed that treatment with sodium acetate or sodium propionate resulted in distinct gene expression changes, while treatment with sodium butyrate mimicked the treatment with a combination of all three SCFAs **(Figure 3a-b)**. Since the different combination treatments greatly overlapped, we focused on the 5 mM combination treatment moving forward, since this treatment had equimolar total concentrations of SCFAs compared to the individual treatments. Analyzing the overlap of differential genes changing with individual SCFAs and the 5 mM SCFA mixture revealed largely overlapping gene changes, with butyrate treatment alone almost completely overlapping with the combination of SCFAs (**Figure 3c-e**). This suggests that the effects of the combination SCFA treatment on gene expression may primarily be due to the action of butyrate.

**Figure 3:**
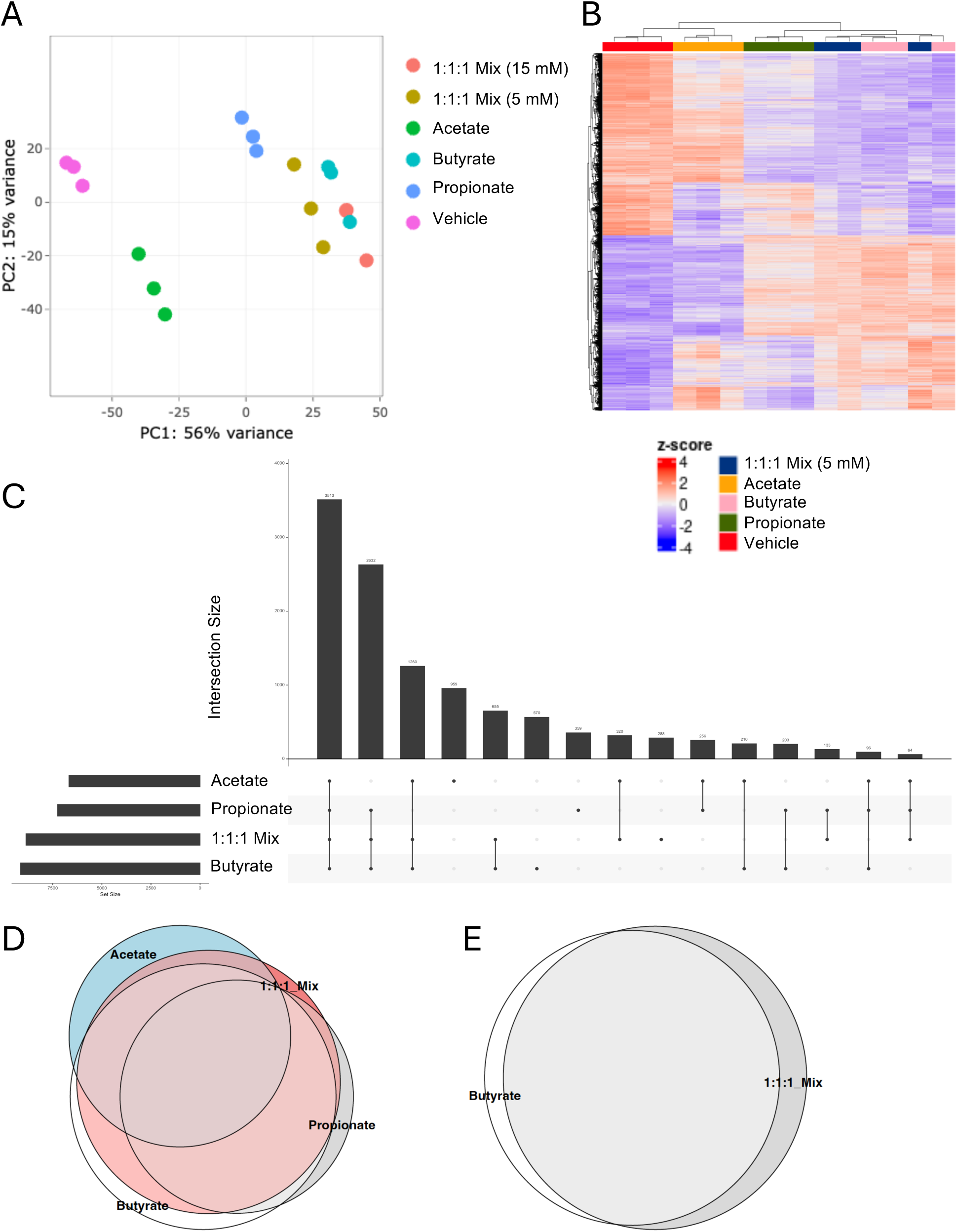
SCFA treatments result in overlapping and distinct gene expression changes. RNA-seq was performed on HCT-116 cells grown in DMEM and treated with 5 mM of single SCFAs or an equimolar combination (1.67 mM) each of acetate, butyrate, and propionate for 6 hours. (A) Principle Component Analysis depicting the variance across all groups. (B) Heatmap of differential gene expression. Differential genes were identified using DESeq2 Wald test and statistically significant genes were defined using a padj < 0.05 cutoff. Displayed are all significant genes after a pairwise comparison between vehicle treatment and the combination SCFA treatment. Genes and samples are hierarchically clustered and values shown are z-scores of regularized log of counts across genes. (C) UpSet plot showing the overlap between significant differential gene expression of different treatment groups compared to vehicle (D-E) Euler diagrams showing the overlap between significant differential gene expression of different treatment groups compared to vehicle.

We also performed RNA-seq analysis with the individual or combination SCFAs in the context of HPLM media (**Supplemental Figure 3**). Here, we analyzed differential gene expression after 24 hours of treatment. At this time point, differentially expressed genes following propionate and butyrate treatments were similarly clustered with genes following both the equimolar and physiological combination of SCFAs (**Supplemental Figure 3a-c**). However, despite the time difference and change in media composition, we generally observed many similar genes changing in both experiments following butyrate treatment (**Supplemental Figure 3d**). Again, butyrate treatment alone almost completely overlapped with SCFA combination treatments, either at the 1:1:1 or 3:1:1 ratios of acetate:butyrate:propionate (**Supplemental Figure 3e**).

Since butyrate largely drives gene expression changes with an SCFA mixture, we decided to focus on butyrate for the remainder of our studies. We next aimed to further analyze how butyrate impacts gene expression across different cell lines and species **(Supplemental Figure 4)**. We first compared the effect of butyrate on gene expression across different cell lines: HCT-116 and Caco-2. Butyrate treatment of these two cell lines resulted in both distinct and overlapping differential genes **(Supplemental Figure 4a)**. We also compared butyrate treatment in these cell lines to an *in vivo* setting: tributyrin gavage in mice, in which we observed dynamic regulation of histone butyrylation in intestinal epithelial cells **(Supplemental Figure 4b)**^11^. We identified 560 genes that overlapped between all three model systems **(Supplemental Figure 4c)**.

These overlapping genes were enriched in cell signaling responses and lipid catabolism **(Supplemental Figure 4d)**. This suggests that butyrate treatment has some common effects on gene expression across cell lines and species.

Since butyrate can have different fates in the intestinal epithelium, we next wanted to define potential chromatin effects with a focus on histone butyrylation. We therefore set up a series of different treatments **(Figure 4a)**. We first wanted to eliminate genes that were changing simply as a result of HDAC inhibition, since butyrate can function as an HDAC inhibitor^9^. We treated cells with another HDAC inhibitor, Entinostat, to identify genes that were targets of HDAC inhibition (HDACi)^25^. We also aimed to target the enzyme that deposits histone butyrylation, which was previously described to be p300/CBP^14,26^. We therefore treated cells with A485, a selective p300/CBP inhibitor, with or without butyrate^27^. Treatment with either inhibitor impacted cell number after 48 or 72 hours, yet had minimal effects at 24 hours **(Supplemental Figure 5)**. Importantly, both inhibitors regulated histone butyrylation and acetylation at H3K27 **(Figure 4b-c).** Since p300/CBP catalyzes H3K27bu as well as acetylation, we reasoned that differential genes changing with A485 treatment that were separate from Entinostat-dependent genes would potentially identify genes that were related to the function of butyrate on chromatin, such as histone butyrylation.

**Figure 4:**
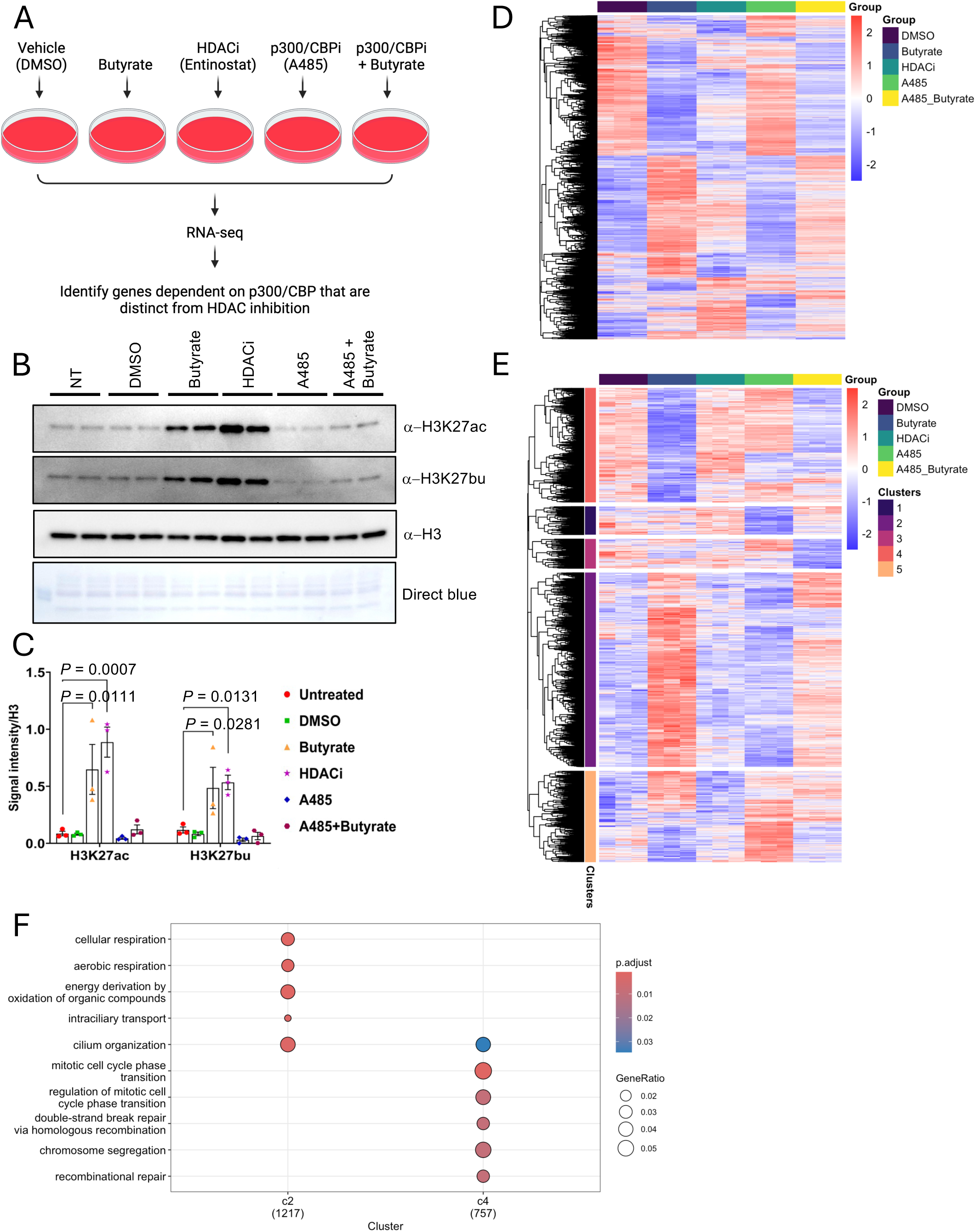
Butyrate treatment has effects independent of HDAC inhibition that rely on p300/CBP activity. (A) Schematic of experimental set-up. Cells were treated with either vehicle (DMSO), 5 mM butyrate, 1 μM Entinostat, 1 μM A485, or A485 with butyrate for 24 hours in HPLM media. Image was created with Biorender.com. (B) Levels of select histone marks were analyzed by immunoblotting with the different treatments. H3 serves as a loading control and representative blot is shown. (C) Quantitative analysis of signal intensity for the histone marks normalized to total histone H3 levels. Immunoblot signal intensities were quantified using Image Lab 6.1, and the bar graphs represent the mean and s.e.m. for three independent experiments (n = 3). Statistical significance was evaluated using one-way ANOVA with Dunnett’s test for multiple comparisons. (D) Heatmap of differential gene expression. Genes were subset to those that are significantly variable across the dataset using Likelihood Ratio Test (padj < 0.05) in DESeq2. Z-scores of regularized log counts across genes are shown. Genes and samples are hierarchically clustered. (E) Heatmap and hierarchical clustering of differential gene expression after removing Entinostat-dependent genes. Entinostat-dependent genes were identified using DESeq2 Wald test and defined using a padj < 0.05 cutoff in DMSO vs HDACi. Genes were subset to those that are significantly variable across the dataset using Likelihood Ratio Test (padj < 0.05) in DESeq2. Z-scores of regularized log of counts across genes are shown. Genes and samples are hierarchically clustered, and tree was cut into 5 separate clusters. (F) Gene Ontology analysis of p300/CBP dependent genes that are independent of HDAC inhibition. GO term enrichment of genes in clusters 2 and 4 from Figure 4e are shown using over representation analysis with Fisher exact test from clusterProfiler.

We observed robust differential gene expression changes following our inhibitor treatments **(Figure 4d)**. Expectedly, butyrate and Entinostat had many differential genes that overlapped, which we attributed to their common HDAC inhibitory roles. In addition, A485 treatment in the presence of butyrate impacted the expression of many genes, the majority of which overlapped with Entinostat **(Figure 4d)**. Next, after removing differential genes changing with Entinostat, we performed hierarchical clustering to identify genes that were differentially regulated by A485 **(Figure 4e).** These genes we identified as p300/CBP dependent, but independent of HDAC inhibition. We performed GO analysis on all clusters, and two subsets (clusters 2 and 4) were significantly enriched for distinct gene categories. We observed genes related to cellular respiration and cell transport were decreased following A485 treatment, while genes related to cell cycle and DNA repair were increased **(Figure 4f)**. This suggests that butyrate has distinct effects on gene regulation that are dependent on p300/CBP activity, but independent of its function as an HDAC inhibitor.

## Discussion

Our study demonstrates that select histone acylations originally found in the mouse intestinal epithelium are also dynamically regulated by metabolites in human colon cancer cells **(Figure 1)**. Consistent with other studies, our work further demonstrates that these histone acylation marks are associated with open chromatin regions **(Figure 2)**. In addition, we detect a representative histone acylation mark, H3K27bu, in human colon samples. This suggests that similar to the *in vivo* setting of the mouse intestine, human samples and cell culture models also harbor unique histone acyl marks that can be regulated by metabolism and metabolite availability.

In the intestinal lumen, SCFAs are often detected as a mixture of acetate, butyrate, and propionate. We aimed to compare the effects of single SCFA treatments vs. mixtures on gene regulation. Surprisingly, our data suggests that in the presence of acetate, butyrate, and propionate, differential genes changing with butyrate completely overlap with the differential genes changing with a mixture of SCFAs **(Figure 3)**. This suggests that butyrate drives effects on gene expression, especially at an earlier time point after treatment. Furthermore, we observe a subset of butyrate-dependent genes that are consistent across different growth conditions, as well as across cell lines and species.

While these associative studies demonstrate that butyrate regulates gene expression, the action of SCFAs can occur through a variety of mechanisms. Here, we aimed to focus on potential chromatin functions of these histone acylations and their impact on gene regulation. The role of butyrate as an HDAC inhibitor has been extensively studied, yet potential functions of histone butyrylation are less known. By using inhibitors to target different fates of butyrate, we could elucidate HDAC inhibitor-independent effects that are governed by the histone acyltransferases p300/CBP **(Figure 4)**. We reasoned that these genes are likely to be connected to histone butyrylation. Importantly, we identified a select gene set that fits this profile as potential histone butyrylation targets, and found genes enriched in chromatin remodeling, transcription, and translation.

The fact that single treatments of butyrate or propionate increase levels of multiple histone acyl marks including acetylation, butyrylation, and propionylation **(Figures 1C-D, Supplemental Figure 2A)** suggests action of these metabolites beyond functioning as donor molecules. These metabolites have additional regulatory effects on chromatin through inhibiting histone deacetylase enzymes, which we speculate is a likely mechanism for these increased acyl marks. Furthermore, this suggests that endogenous levels of histone butyrylation and propionylation can be uncovered through HDAC inhibition, which is supported both by the inhibitory action of these metabolites and by the detection of these marks in untreated cells^9,14,26,28^.

A class I HDAC inhibitor, Entinostat, also increases levels of histone butyrylation analogous to acetylation **(Figure 4B)**. This observation suggests that histone butyrylation may be regulated by the activity of class I HDACs. Interestingly, histone butyrylation is thought to generally be erased by Sirtuins, or class III HDACs^29^.

However, histone crotonylation, which is the same length as histone butyrylation but with a double bond, can be regulated by both sirtuins and class I HDACs^30,31^. How different classes of erasers function to remove histone butyrylation and potential side chain differences in the dynamics of histone acylations are important open questions for future investigation.

## Methods

### Cell Culture & Treatments

HCT-116 and Caco-2 cells were obtained from ATCC. HCT-116 cells were grown in DMEM medium (Gibco 11965-092) with 10% fetal bovine serum (FBS, Gibco A5669701) and 1% penicillin/streptomycin (Gibco 15140-122). Caco-2 cells were grown in Eagle’s Minimum Essential Medium (MEM medium (Gibco 10370-021) supplemented with 2 mM GlutaMAX, 1 mM sodium pyruvate, 20% FBS, and 1% penicillin/streptomycin). When cells were grown in human plasma like media (HPLM, Gibco A4899101), cells were washed in PBS and cultured in HPLM for at least three days prior to experiments. All cell lines were regularly tested for Mycoplasma (ABM, G238) and confirmed with STR testing (ATCC). Cells were grown in conditions of 37 °C and 5% CO_2_. For SCFA treatments, cells were treated with 5 mM of metabolites for 24 hours, unless specified differently in the figure legends. Metabolites were dissolved in sterile water and used are as follows: sodium acetate (Sigma-Aldrich S1429), sodium butyrate (Sigma-Aldrich 1.37127.0250), and sodium propionate (Sigma-Aldrich 18108).

### Histone Extraction

Cells were harvested and lysed in hypotonic buffer. The supernatant was saved as cell lysate, and the pellet was processed for acid extraction of histones^32^. Briefly, pellets were vortexed in 0.4 N sulfuric acid and incubated overnight, then precipitated with 100% trichloroacetic acid. After precipitation, pellets were washed with acetone and allowed to air dry. Histones were then resuspended in water and protein concentration was assessed with BCA assay (Thermo 23225) and Coomassie staining.

### Immunoblotting

Samples were run on 16% or 4-20% tris glycine SDS-PAGE gels (Invitrogen XP00165BOX or XP04205BOX) and transferred to PVDF membranes. Blocking and all antibody incubations were done in 5% milk, with TBST washes in between.

Immunoblots were imaged using Immobilon ECL (Millipore WBKLS0500) and an ChemiDoc MP Imaging System (BioRad). Antibodies used are as follows: H3K27bu (Millipore ABE2854, 1:2,000), H3K9bu (PTM-Biolabs PTM-305, 1:5,000), H3K27ac (Active Motif 39133, 1:5,000), H3K9ac (Invitrogen MA5-33384, 1:5,000), and histone H3 (Abcam, ab1791, 1:100,000).

### Immunofluorescence

Paraffin-embedded human benign intestinal tissue microarrays were received from HistoWiz and stained as previously described^11^. Briefly, sections were rehydrated, boiled in sodium citrate buffer, pH 6.0 (10 mM sodium citrate, 0.05% Tween 20), and then stained with the following antibodies or stains: a-H3K27bu (Millipore ABE2854, 1:500), a-Villin-647 (SantaCruz sc58897-647 lot D2920, 1:200), Goat anti-Rabbit Alexa Fluor 488 (Invitrogen A-11008, 1:1,000), DAPI (1:1,000). Imaging was performed on a Zeiss Inverted LSM 780 laser scanning confocal microscope using a 40x objective.

### GLO Assay

Approximately 1,000 HCT-116 cells in HPLM culture medium were seeded per well in three 96-well plates to evaluate cell viability at three time points (24 h, 48 h, and 72 h). The plates were incubated at 37°C with 5% CO₂. On the following day, the culture medium was replaced with treatment medium containing: 5 mM of individual SCFAs; 1 µM of A485, a p300/CBP inhibitor (MedChemExpress, HY-107455), a combination of 1 µM A485 and 5 mM butyrate; or 1 µM of Entinostat (ES), a class I HDAC inhibitor (MedChemExpress, HY-12163). Cells treated with DMSO and 5 mM NaCl served as vehicle controls. Treatments were performed in triplicate for each condition. After 24 hours, cell viability for one 96-well plate was measured by adding an equal volume of CellTiter-Glo® Reagent to the cell culture medium in each well. The contents were mixed on an orbital shaker for 2 minutes to induce cell lysis, incubated at room temperature for 10 minutes, and luminescence was recorded using the VICTOR Nivo™ multimode plate reader. For the remaining two plates, treatment medium was replaced with regular culture media after 24 hours, and cell viability was assessed at 48- and 72-hours post-treatment.

### RNA Isolation

Total RNA was isolated from cells using the RNeasy Mini Kit (Qiagen 74104) or Quick-RNA Miniprep Kit (Zymo R1054) with on-column DNA digestion. RNA samples were analyzed on the Bioanalyzer RNA 6000 Pico (Agilent) or TapeStation prior to library preparation.

### Chromatin Immunoprecipitation

Cells were fixed in PBS with 1% formaldehyde for 10 minutes, followed by quenching with 125 mM glycine for 5 minutes. Crosslinked cells were washed once with PBS and then lysed following the NEXSON protocol^33^. Briefly, cells were resuspended in FL Buffer (5 mM PIPES, pH 8.0, 85 mM KCl, 0.5% NP-40, protease inhibitor cocktail) and then sonicated in the Bioruptor Pico (Diagenode) on low power at 15 seconds on 30 seconds off cycles to lyse cell membranes and release nuclei. Nuclei were pelleted, washed in FL Buffer, and then resuspended in D3 Sonication Buffer (10 mM Tris-HCl, pH 8.0, 0.1% SDS, 1 mM EDTA, protease inhibitor cocktail) Sonication was performed in the Covaris E220 (Covaris) and checked prior to proceeding with immunoprecipitations (IPs). For IPs, 30 μg sonicated chromatin was diluted into ChIP Dilution Buffer (10 mM Tris-HCl, pH 8.0, 1 mM EDTA, 150 mM NaCl, 1% Triton-X, protease inhibitor cocktail) and added to antibody-bound Protein A Dynabeads (Invitrogen 10002D) and incubated overnight in the cold room, rotating. Antibodies used were as follows: H3K4me3 (Active Motif 39159), H3K27bu (Millipore ABE2854), H3K9bu (PTM Biolabs PTM-305), H3K27pr (Millipore, ABE2852), H3K9pr (Millipore, ABE2852). The next day, beads were washed six times with RIPA buffer (50 mM HEPES-KOH, pH 7.5, 100 mM LiCl, 1 mM EDTA, 0.7% Na-Deoxycholate, 1% NP-40) followed by one wash with TE-NaCl Buffer (10 mM Tris-HCl, pH 8.0, 50 mM NaCl, 1 mM EDTA). DNA was eluted from the beads using Elution Buffer (50 mM Tris-HCl, pH 8.0, 10 mM EDTA, 1% SDS). DNA was reverse crosslinked at 65 °C overnight, followed by RNase and ProteinaseK digestion and column purification using the ChIP DNA Clean & Concentrator kit (Zymo D5205). DNA was quantified using the Qubit 4 Fluorometer (Thermo).

### Library preparations & sequencing

RNA libraries were prepared using the NEBNext Poly(A) mRNA Magnetic Isolation Module (NEB E7490L) and NEBNext Ultra II RNA Library Prep Kit for Illumina (NEB E7770L). ChIP libraries were prepared using NEBNext Ultra II DNA Library Prep Kit for Illumina (NEB E7645L). All kits were used according to the manufacturers instructions. All libraries were analyzed for quality by TapeStation prior to sequencing. Single-end sequencing was performed on the Illumina NextSeq 500 sequencer and paired-end sequencing was performed at Admera Health, LLC. Paired-end data was processed as single-end to be consistent across experiments.

### Bioinformatics Analysis

Sequence and transcript coordinates for human hg19 UCSC genome and gene models were retrieved from knownGene (v.3.4.0) Bioconductor libraries: BSgenome.Hsapiens.UCSC.hg19 and TxDb.Hsapiens.UCSC.hg19.knownGene. For the analysis of RNA-seq data, transcript expressions were calculated using Salmon quantification software^34^ (v.0.8.2) and gene expression levels as TPMs and counts were retrieved using Tximport^35^ (v.1.8.0). Normalization and rlog transformation of raw read counts in genes, principal-component analysis and differential gene expression analysis were performed using DESeq2 (v.1.20.0). To visualize overlapping differential genes, the UpSetR package (v.1.4.0) and eulerr package (7.0.2) were used^36^. Heatmaps were made with pheatmap (v 1.0.12) based on subsets of genes derived from DESeq2 analysis (indicated in figure legends). For hierarchical clustering, significantly variable genes were determined using a Likelihood Ratio Test (padj < 0.05). The z score of the rlog of gene counts was used as the input for clustering. Hierarchical clustering was performed using pheatmap. Gene set enrichment tests were conducted using clusterProfiler^37^ (v.3.18.1). GO analysis was performed using over representation analysis and the Fisher exact test in DAVID^38,39^. For the analysis of ChIP-seq data, reads were mapped using the Rsubread package’s align function^40^ (v.1.30.6). Genome distribution was determined using the ChIPQC package (v.1.16.2)^41^. Heat maps were generated with deeptools^42^ (v.3.5). Normalized, fragment-extended signal bigWigs were created using the rtracklayer package^43^ (v.1.40.6) and then visualized and exported from IGV.

### Statistics

Details for statistical tests and replicates are described in the figure legends. All measurements shown were taken from distinct biological samples unless otherwise indicated. No data points were excluded from analysis. Data distribution was assumed to be normal but this was not formally tested, unless otherwise noted. Prism 10 was used to generate plots and perform statistical tests. Error bars represent the standard error. Unpaired two-tailed Student’s t-test, one-way ANOVA with multiple comparisons, or Likelihood Ratio Test were used to assess significance and is indicated in the figure legends. P<0.05 was considered statistically significant.

## Data Availability

The RNA-seq and ChIP-seq data have been deposited to the NCBI GEO under accession number GSE298547. Any additional data that support the findings of this study or materials are available from the corresponding author upon request.

## Code Availability

No custom code was used for this study.

## Acknowledgements

We thank all members of the former C. David Allis laboratory where this study originated for helpful discussions and support. This article is dedicated to the memory of C. David Allis, who died on January 8, 2023. We acknowledge the help and support of the Genomics, Bioimaging, and Bioinformatics Resource Centers at the Rockefeller University. Select schematics were generated using BioRender. This work was supported by the National Institutes of Health (R00GM143550 to L.A.G.) and start-up funds from the Department of Biochemistry at Case Western Reserve University School of Medicine.

## Author Contributions

L.A.G. conceptualized this project. L.A.G., T.K., and C.A.C. wrote the manuscript with input from all authors. L.A.G., T.K., C.A.C., Z.K.L., M.R.S., A.M.D., Z.N., and M.R.P. conducted experiments, contributed to sample preparation, and provided conceptual advice. T.S.C. and L.A.G. participated in study supervision.

---

**Supplemental Figure 1:**
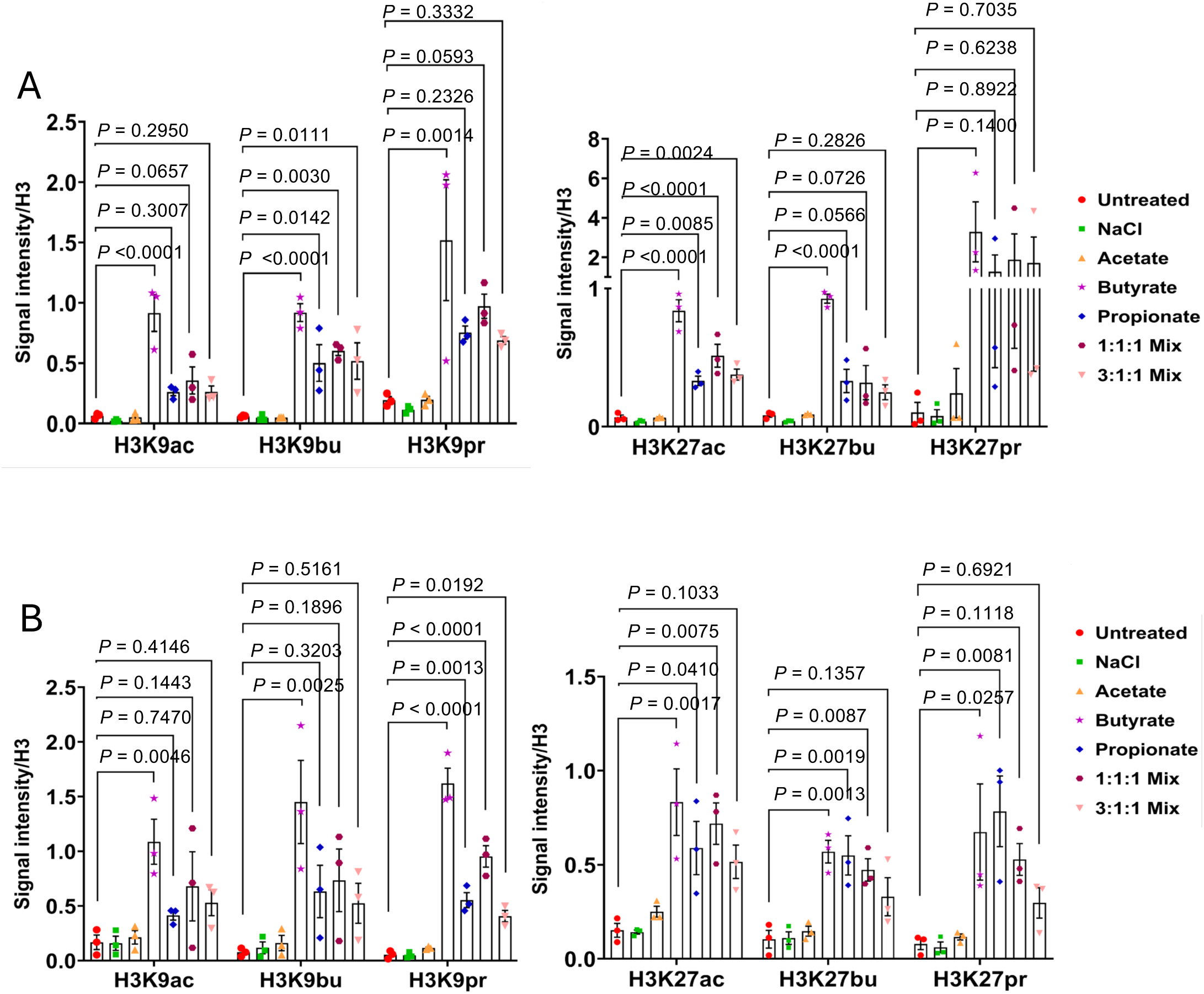
Quantification of immunoblotting data related to Figure 1. Histones were extracted from (A) HCT116 cells cultured in HPLM medium or (B) Caco-2 cells and treated for 24 hours with 5 mM of the following treatments: NaCl, acetate, butyrate, propionate, 1:1:1 Mix (equimolar SCFA mixture, 1.67 mM each), and 3:1:1 Mic (physiological SCFA ratios: 3 mM acetate, 1 mM butyrate, and 1 mM propionate). Quantitative analysis of signal intensity for the histone marks normalized to total histone H3 levels. Immunoblot signal intensities were quantified using Image Lab 6.1, and the bar graphs represent the mean and s.e.m. for three independent experiments (n = 3). Statistical significance was evaluated using one-way ANOVA with Dunnett’s test for multiple comparisons.

**Supplemental Figure 2:**
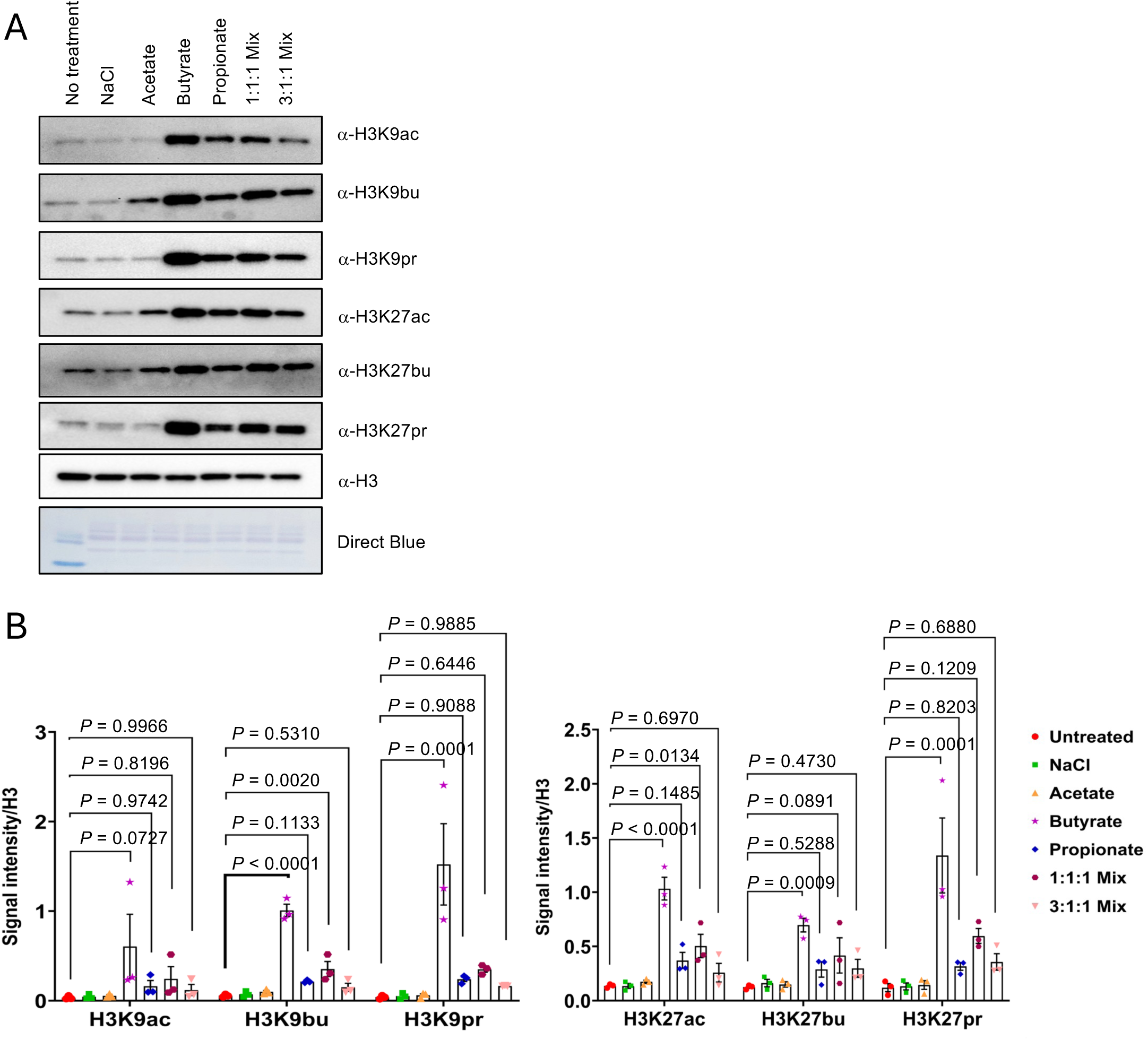
Immunoblotting data in HCT-116 cells grown in DMEM. HCT116 cells cultured in DMEM medium were treated for 24 hours with 5 mM of the following treatments: NaCl, acetate, butyrate, propionate, 1:1:1 Mix (equimolar SCFA mixture, 1.67 mM each), and 3:1:1 Mix (physiological SCFA ratios: 3 mM acetate, 1 mM butyrate, and 1 mM propionate). (A) Representative immunoblot of n =3 independent biological replicates. NaCl serves as a control for sodium addition and H3 serves as a loading control. (B) Quantitative analysis of signal intensity for the histone marks normalized to total histone H3 levels. Immunoblot signal intensities were quantified using Image Lab 6.1, and the bar graphs represent the mean and s.e.m. for three independent experiments (n = 3). Statistical significance was evaluated using one-way ANOVA with Dunnett’s test for multiple comparisons.

**Supplemental Figure 3:**
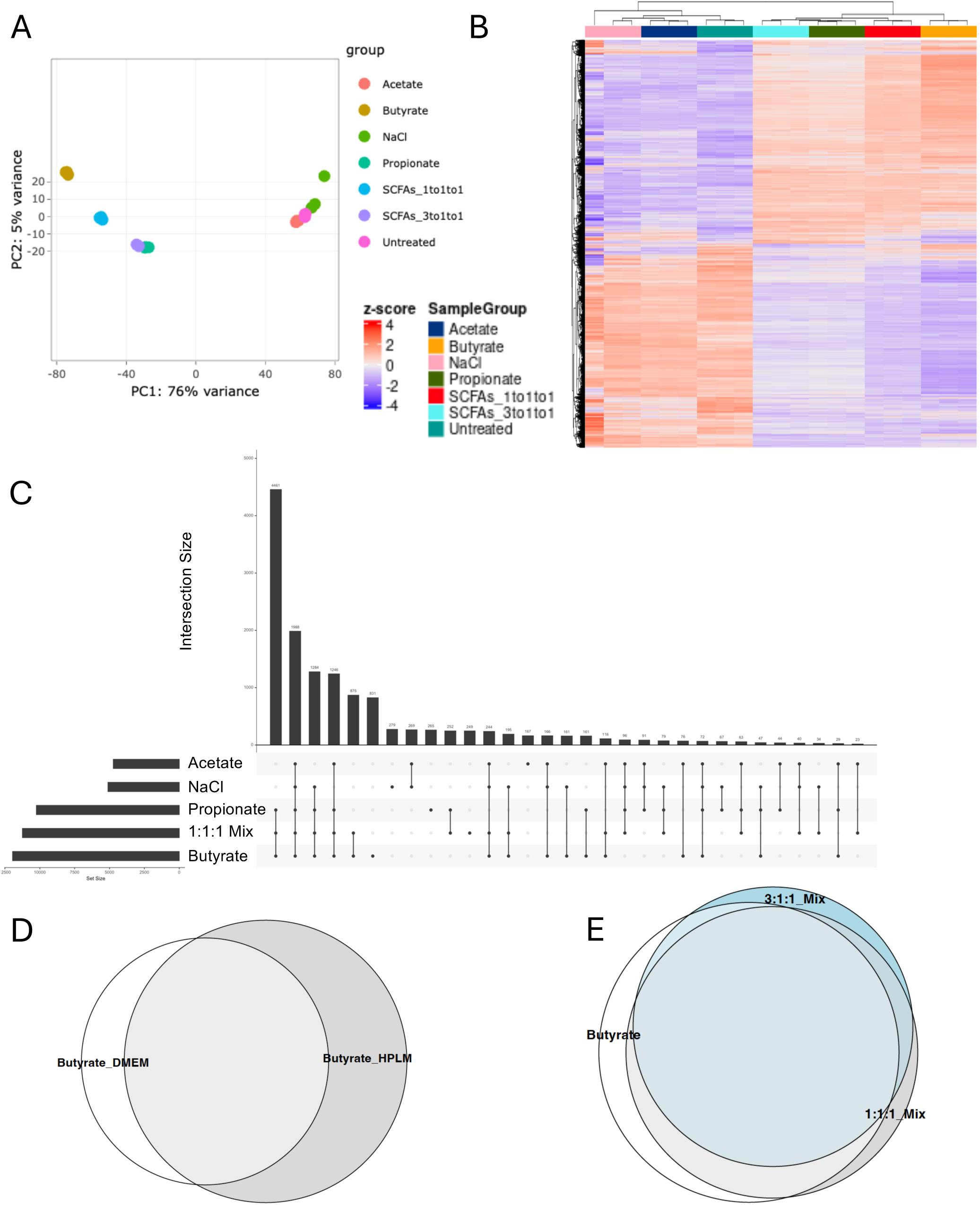
SCFA treatment in HPLM generally recapitulates gene expression changes observed in DMEM. RNA-seq was performed on cells treated with a sodium matched control (NaCl), 5 mM of single SCFAs, an equimolar combination (1.67 mM) each of acetate, butyrate, and propionate (1:1:1 mix), or physiological combination (3:1:1 mix of 3 mM acetate, 1 mM butyrate, 1 mM propionate) for 24 hours. (A) Principal component analysis of all samples. (B) Heatmap of differential gene expression. Differential genes were identified using DESeq2 Wald test and statistically significant genes were defined using a padj < 0.05 cutoff. Displayed are all significant genes changing in the 3:1:1 SCFA mixture versus untreated samples. Genes and samples are hierarchically clustered and values shown are z-scores of regularized log counts across genes. (C) UpSet plot showing the overlap between significant differential gene expression of different treatment groups compared to vehicle (D-E) Euler diagrams showing the overlap of significant differential genes compared to untreated cells (D) following butyrate treatments across experiments and (E) following butyrate or SCFA mixes in HPLM.

**Supplemental Figure 4:**
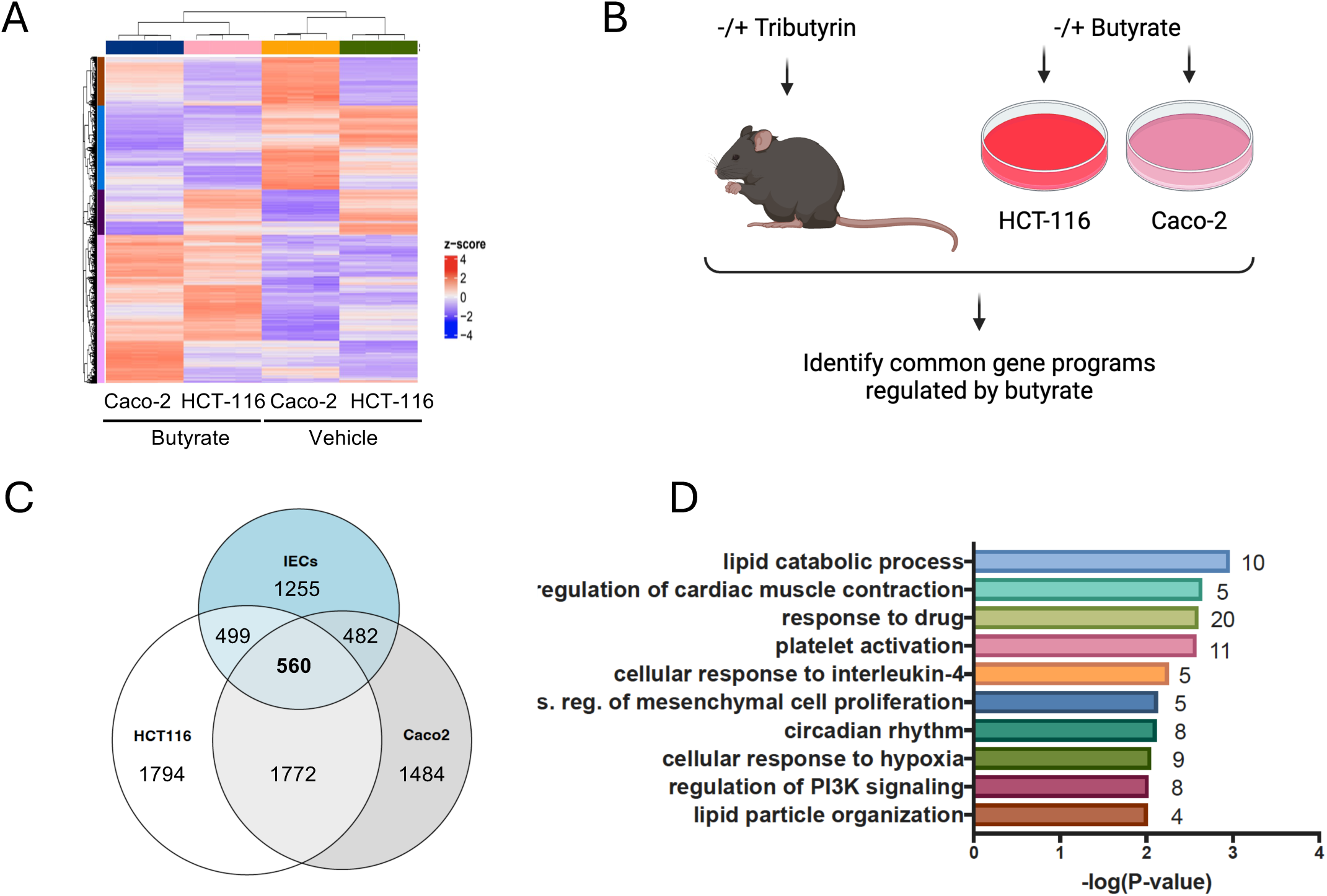
Comparison of gene expression changes from butyrate treatments in different human cell lines and mouse intestinal epithelial cells. (A) Heatmap showing differential gene expression in different human cell lines, HCT-116 and Caco-2, in response to butyrate treatment. Differential genes were identified using the DESeq2 package and defined using a padj < 0.05 and log2fc > 2 or < −2 as cut offs. Genes and samples are hierarchically clustered and values shown are z-scores of regularized log counts across genes. (B) Schematic showing different treatments in mice and human cell lines, generated in Biorender.com. (C). Euler diagram showing overlap between human cell lines and mouse IECs. Mouse IEC genes are defined as all significant differentially expressed genes from tributyrin treatment with ampicillin compared to ampicillin treatment and mock gavage from GSE216319. Human cell lines genes are defined as significantly changed genes 2< fold change >2. (D) Gene ontology categories for the 560 common genes from mouse IECs and both cell lines treated with butyrate. GO analysis was performed using over representation analysis and the Fisher exact test in DAVID. The top 10 Biological Process categories are shown.

**Supplemental Figure 5:**
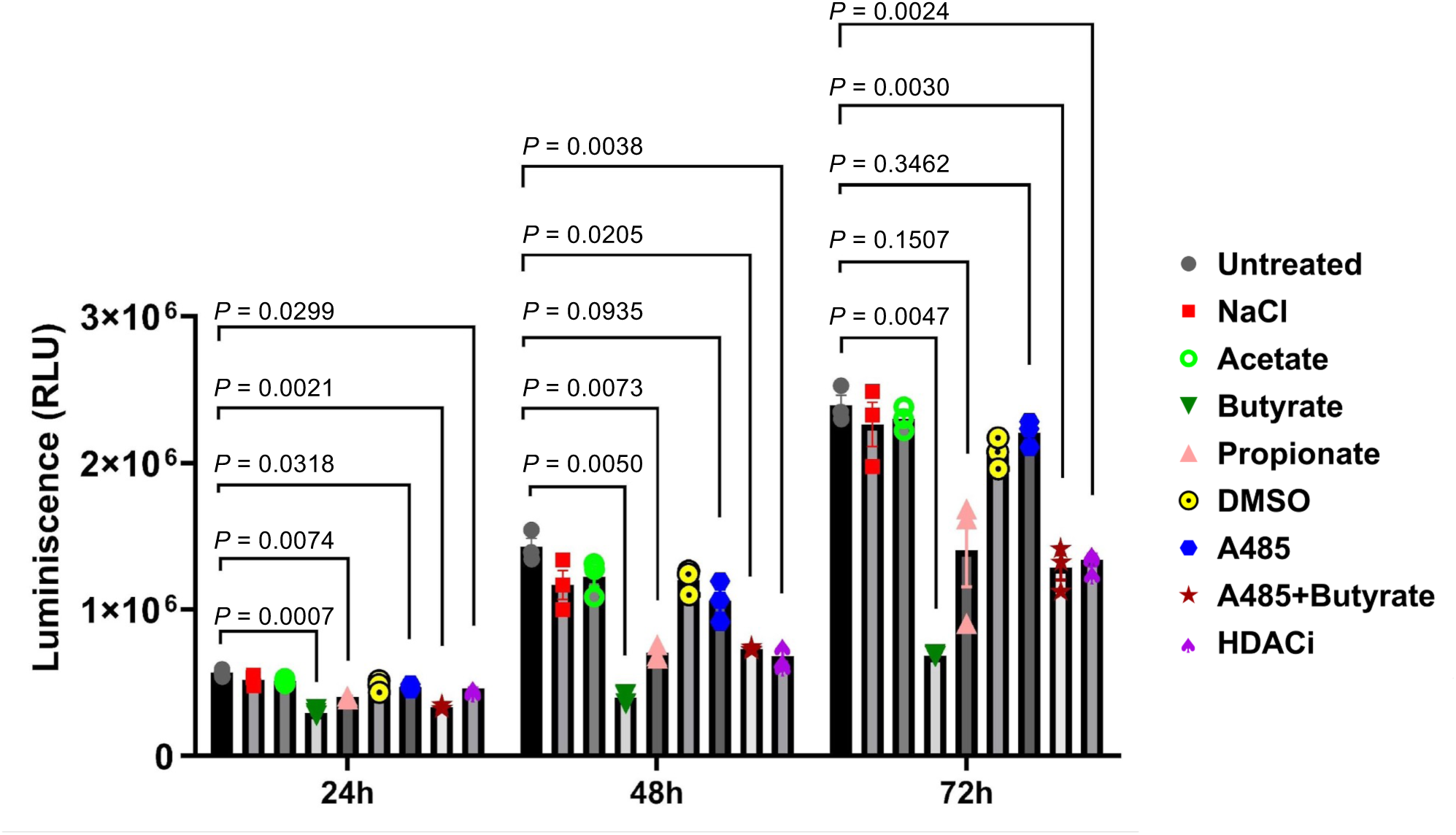
Cell viability following treatments with SCFAs, a p300/CBP inhibitor, and an HDAC inhibitor. HCT-116 cells in HPLM culture medium were treated with media containing: 5 mM of SCFAs; 1 µM of A485 (a p300/CBP inhibitor); a combination of 1 µM A485 and 5 mM butyrate; or 1 µM of Entinostat (ES), a class I HDAC inhibitor. Cells treated with DMSO served as vehicle controls. Treatments were performed in triplicate for each condition. Cell viability was measured by CellTiter-Glo®. Results are presented as the mean and s.e.m. (n = 3). Statistical analysis was performed using two-way ANOVA followed by Dunnett’s test for multiple comparisons.

## References

1. Cummings, J. H., Pomare, E. W., Branch, W. J., Naylor, C. P. & Macfarlane, G. T. Short chain fatty acids in human large intestine, portal, hepatic and venous blood. Gut 28, 1221–7 (1987).

2. Besten, G. den, Eunen, K. van, Groen, A. K., Venema, K., Reijngoud, D.-J. & Bakker, B. M. The role of short-chain fatty acids in the interplay between diet, gut microbiota, and host energy metabolism. J Lipid Res 54, 2325–40 (2013).

3. Koh, A., De Vadder, F., Kovatcheva-Datchary, P. & Bäckhed, F. From Dietary Fiber to Host Physiology: Short-Chain Fatty Acids as Key Bacterial Metabolites. Cell 165, 1332–1345 (2016).

4. Fleming, S. E., Fitch, M. D., DeVries, S., Liu, M. L. & Kight, C. Nutrient utilization by cells isolated from rat jejunum, cecum and colon. J Nutr 121, 869–78 (1991).

5. Roediger, W. E. Utilization of nutrients by isolated epithelial cells of the rat colon. Gastroenterology 83, 424–9 (1982).

6. Brown, A. J., Goldsworthy, S. M., Barnes, A. A., Eilert, M. M., Tcheang, L., Daniels, D., Muir, A. I., Wigglesworth, M. J., Kinghorn, I., Fraser, N. J., Pike, N. B., Strum, J. C., Steplewski, K. M., Murdock, P. R., Holder, J. C., Marshall, F. H., Szekeres, P. G., Wilson, S., Ignar, D. M., Foord, S. M., Wise, A. & Dowell, S. J. The Orphan G protein-coupled receptors GPR41 and GPR43 are activated by propionate and other short chain carboxylic acids. J Biol Chem 278, 11312–9 (2003).

7. Poul, E. L., Loison, C., Struyf, S., Springael, J.-Y., Lannoy, V., Decobecq, M.-E., Brezillon, S., Dupriez, V., Vassart, G., Damme, J. V., Parmentier, M. & Detheux, M. Functional characterization of human receptors for short chain fatty acids and their role in polymorphonuclear cell activation. J Biol Chem 278, 25481–9 (2003).

8. Donohoe, D. R., Collins, L. B., Wali, A., Bigler, R., Sun, W. & Bultman, S. J. The Warburg effect dictates the mechanism of butyrate-mediated histone acetylation and cell proliferation. Mol Cell 48, 612–26 (2012).

9. Vidali, G., Boffa, L. C., Bradbury, E. M. & Allfrey, V. G. Butyrate suppression of histone deacetylation leads to accumulation of multiacetylated forms of histones H3 and H4 and increased DNase I sensitivity of the associated DNA sequences. Proc Natl Acad Sci U S A 75, 2239–43 (1978).

10. Krautkramer, K. A., Kreznar, J. H., Romano, K. A., Vivas, E. I., Barrett-Wilt, G. A., Rabaglia, M. E., Keller, M. P., Attie, A. D., Rey, F. E. & Denu, J. M. Diet-Microbiota Interactions Mediate Global Epigenetic Programming in Multiple Host Tissues. Mol. Cell 64, 982–992 (2016).

11. Gates, L. A., Reis, B. S., Lund, P. J., Paul, M. R., Leboeuf, M., Djomo, A. M., Nadeem, Z., Lopes, M., Vitorino, F. N., Unlu, G., Carroll, T. S., Birsoy, K., Garcia, B. A., Mucida, D. & Allis, C. D. Histone butyrylation in the mouse intestine is mediated by the microbiota and associated with regulation of gene expression. Nat. Metab. 6, 697–707 (2024).

12. Allis, C. D. & Jenuwein, T. The molecular hallmarks of epigenetic control. Nature Genetics 17, 487–500 (2016).

13. Gates, L. A., Foulds, C. E. & O’Malley, B. W. Histone Marks in the ‘Driver’s Seat’: Functional Roles in Steering the Transcription Cycle. Trends Biochem. Sci. 42, 977–989 (2017).

14. Tan, M., Luo, H., Lee, S., Jin, F., Yang, J. S., Montellier, E., Buchou, T., Cheng, Z., Rousseaux, S., Rajagopal, N., Lu, Z., Ye, Z., Zhu, Q., Wysocka, J., Ye, Y., Khochbin, S., Ren, B. & Zhao, Y. Identification of 67 histone marks and histone lysine crotonylation as a new type of histone modification. Cell 146, 1016–28 (2011).

15. Wellen, K. E., Hatzivassiliou, G., Sachdeva, U. M., Bui, T. V., Cross, J. R. & Thompson, C. B. ATP-citrate lyase links cellular metabolism to histone acetylation. Science 324, 1076–80 (2009).

16. Berger, S. L. & Sassone-Corsi, P. Metabolic Signaling to Chromatin. Cold Spring Harb Perspect Biol 8, (2016).

17. Sabari, B. R., Zhang, D., Allis, C. D. & Zhao, Y. Metabolic regulation of gene expression through histone acylations. Nat. Rev. Mol. Cell Biol. 18, 90–101 (2017).

18. Dai, Z., Ramesh, V. & Locasale, J. W. The evolving metabolic landscape of chromatin biology and epigenetics. Nat. Rev. Genet. 21, 737–753 (2020).

19. Simithy, J., Sidoli, S., Yuan, Z.-F., Coradin, M., Bhanu, N. V., Marchione, D. M., Klein, B. J., Bazilevsky, G. A., McCullough, C. E., Magin, R. S., Kutateladze, T. G., Snyder, N. W., Marmorstein, R. & Garcia, B. A. Characterization of histone acylations links chromatin modifications with metabolism. Nat. Commun. 8, 1141 (2017).

20. Lund, P. J., Gates, L. A., Leboeuf, M., Smith, S. A., Chau, L., Lopes, M., Friedman, E. S., Saiman, Y., Kim, M. S., Shoffler, C. A., Petucci, C., Allis, C. D., Wu, G. D. & Garcia, B. A. Stable isotope tracing in vivo reveals a metabolic bridge linking the microbiota to host histone acetylation. Cell Rep. 41, 111809 (2022).

21. Kebede, A. F., Nieborak, A., Shahidian, L. Z., Gras, S. L., Richter, F., Gómez, D. A., Baltissen, M. P., Meszaros, G., Magliarelli, H. de F., Taudt, A., Margueron, R., Colomé-Tatché, M., Ricci, R., Daujat, S., Vermeulen, M., Mittler, G. & Schneider, R. Histone propionylation is a mark of active chromatin. Nat Struct Mol Biol 24, 1048–1056 (2017).

22. Goudarzi, A., Zhang, D., Huang, H., Barral, S., Kwon, O. K., Qi, S., Tang, Z., Buchou, T., Vitte, A.-L., He, T., Cheng, Z., Montellier, E., Gaucher, J., Curtet, S., Debernardi, A., Charbonnier, G., Puthier, D., Petosa, C., Panne, D., Rousseaux, S., Roeder, R. G., Zhao, Y. & Khochbin, S. Dynamic Competing Histone H4 K5K8 Acetylation and Butyrylation Are Hallmarks of Highly Active Gene Promoters. Mol Cell 62, 169–180 (2016).

23. Nshanian, M., Gruber, J. J., Geller, B. S., Chleilat, F., Lancaster, S. M., White, S. M., Alexandrova, L., Camarillo, J. M., Kelleher, N. L., Zhao, Y. & Snyder, M. P. Short-chain fatty acid metabolites propionate and butyrate are unique epigenetic regulatory elements linking diet, metabolism and gene expression. Nat Metab 7, 196–211 (2025).

24. Moussaieff, A., Rouleau, M., Kitsberg, D., Cohen, M., Levy, G., Barasch, D., Nemirovski, A., Shen-Orr, S., Laevsky, I., Amit, M., Bomze, D., Elena-Herrmann, B., Scherf, T., Nissim-Rafinia, M., Kempa, S., Itskovitz-Eldor, J., Meshorer, E., Aberdam, D. & Nahmias, Y. Glycolysis-mediated changes in acetyl-CoA and histone acetylation control the early differentiation of embryonic stem cells. Cell Metab 21, 392–402 (2015).

25. Saito, A., Yamashita, T., Mariko, Y., Nosaka, Y., Tsuchiya, K., Ando, T., Suzuki, T., Tsuruo, T. & Nakanishi, O. A synthetic inhibitor of histone deacetylase, MS-27–275, with marked in vivo antitumor activity against human tumors. Proc Natl Acad Sci U S A 96, 4592–7 (1999).

26. Chen, Y., Sprung, R., Tang, Y., Ball, H., Sangras, B., Kim, S. C., Falck, J. R., Peng, J., Gu, W. & Zhao, Y. Lysine Propionylation and Butyrylation Are Novel Post-translational Modifications in Histones. Mol. Cell. Proteom. 6, 812–819 (2007).

27. Lasko, L. M., Jakob, C. G., Edalji, R. P., Qiu, W., Montgomery, D., Digiammarino, E. L., Hansen, T. M., Risi, R. M., Frey, R., Manaves, V., Shaw, B., Algire, M., Hessler, P., Lam, L. T., Uziel, T., Faivre, E., Ferguson, D., Buchanan, F. G., Martin, R. L., Torrent, M., Chiang, G. G., Karukurichi, K., Langston, J. W., Weinert, B. T., Choudhary, C., Vries, P. de, Drie, J. H. V., McElligott, D., Kesicki, E., Marmorstein, R., Sun, C., Cole, P. A., Rosenberg, S. H., Michaelides, M. R., Lai, A. & Bromberg, K. D. Discovery of a selective catalytic p300/CBP inhibitor that targets lineage-specific tumours. Nature 550, 128–132 (2017).

28. Kim, M., Qie, Y., Park, J. & Kim, C. H. Gut Microbial Metabolites Fuel Host Antibody Responses. Cell Host & Microbe 20, 202–214 (2016).

29. Smith, B. C. & Denu, J. M. Acetyl-lysine Analog Peptides as Mechanistic Probes of Protein Deacetylases. Journal of Biological Chemistry 282, 37256–37265 (2007).

30. Wei, W., Liu, X., Chen, J., Gao, S., Lu, L., Zhang, H., Ding, G., Wang, Z., Chen, Z., Shi, T., Li, J., Yu, J. & Wong, J. Class I histone deacetylases are major histone decrotonylases: evidence for critical and broad function of histone crotonylation in transcription. Cell Research 27, 898–915 (2017).

31. Kelly, R. D. W., Chandru, A., Watson, P. J., Song, Y., Blades, M., Robertson, N. S., Jamieson, A. G., Schwabe, J. W. R. & Cowley, S. M. Histone deacetylase (HDAC) 1 and 2 complexes regulate both histone acetylation and crotonylation in vivo. Sci Rep 8, 14690 (2018).

32. Shechter, D., Dormann, H. L., Allis, C. D. & Hake, S. B. Extraction, purification and analysis of histones. Nat. Protoc. 2, 1445–1457 (2007).

33. Arrigoni, L., Richter, A. S., Betancourt, E., Bruder, K., Diehl, S., Manke, T. & Bönisch, U. Standardizing chromatin research: a simple and universal method for ChIP-seq. Nucleic Acids Res. 44, e67–e67 (2016).

34. Patro, R., Duggal, G., Love, M. I., Irizarry, R. A. & Kingsford, C. Salmon provides fast and bias-aware quantification of transcript expression. Nature Methods 14, 417– 419 (2017).

35. Love, M. I., Hogenesch, J. B. & Irizarry, R. A. Modeling of RNA-seq fragment sequence bias reduces systematic errors in transcript abundance estimation. Nature Biotechnology 34, 1287–1291 (2016).

36. Lex, A., Gehlenborg, N., Strobelt, H., Vuillemot, R. & Pfister, H. UpSet: Visualization of Intersecting Sets. IEEE Transactions on Visualization and Computer Graphics 20, 1983–1992 (2014).

37. Yu, G., Wang, L.-G., Han, Y. & He, Q.-Y. clusterProfiler: an R Package for Comparing Biological Themes Among Gene Clusters. OMICS: A Journal of Integrative Biology 16, 284–287 (2012).

38. Huang, D. W., Sherman, B. T. & Lempicki, R. A. Bioinformatics enrichment tools: paths toward the comprehensive functional analysis of large gene lists. Nucleic Acids Res 37, 1–13 (2009).

39. Huang, D. W., Sherman, B. T. & Lempicki, R. A. Systematic and integrative analysis of large gene lists using DAVID bioinformatics resources. Nat Protoc 4, 44–57 (2009).

40. Liao, Y., Smyth, G. K. & Shi, W. The R package Rsubread is easier, faster, cheaper and better for alignment and quantification of RNA sequencing reads. Nucleic Acids Research 47, e47–e47 (2019).

41. Carroll, T. S., Liang, Z., Salama, R., Stark, R. & Santiago, I. de. Impact of artifact removal on ChIP quality metrics in ChIP-seq and ChIP-exo data. Frontiers in Genetics Volume 5-2014, (2014).

42. Ramírez, F., Ryan, D. P., Grüning, B., Bhardwaj, V., Kilpert, F., Richter, A. S., Heyne, S., Dündar, F. & Manke, T. deepTools2: a next generation web server for deep-sequencing data analysis. Nucleic Acids Research 44, W160–W165 (2016).

43. Lawrence, M., Gentleman, R. & Carey, V. rtracklayer: an R package for interfacing with genome browsers. Bioinformatics 25, 1841–1842 (2009).

